# A mouse model of TB-associated lung fibrosis reveals persistent inflammatory macrophage populations during treatment

**DOI:** 10.1101/2024.06.04.597479

**Authors:** Julie Boucau, Threnesan Naidoo, Yuming Liu, Shatavisha Dasgupta, Neha Jain, Jennie Ruelas Castillo, Nicholas E. Jacobson, Kievershen Nargan, Beth A. Cimini, Kevin W. Eliceiri, Adrie J.C. Steyn, Amy K. Barczak

## Abstract

Post-TB lung disease (PTLD) causes a significant burden of global disease. Fibrosis is a central component of many clinical features of PTLD. To date, we have a limited understanding of the mechanisms of TB-associated fibrosis and how these mechanisms are similar to or dissimilar from other fibrotic lung pathologies. We have adapted a mouse model of TB infection to facilitate the mechanistic study of TB-associated lung fibrosis. We find that the morphologies of fibrosis that develop in the mouse model are similar to the morphologies of fibrosis observed in human tissue samples. Using Second Harmonic Generation (SHG) microscopy, we are able to quantify a major component of fibrosis, fibrillar collagen, over time and with treatment. Inflammatory macrophage subpopulations persist during treatment; matrix remodeling enzymes and inflammatory gene signatures remain elevated. Our mouse model suggests that there is a therapeutic window during which adjunctive therapies could change matrix remodeling or inflammatory drivers of tissue pathology to improve functional outcomes after treatment for TB infection.

## Introduction

Tuberculosis (TB) is the leading cause of death from infection globally^1^; an estimated 10 million individuals were treated for active TB disease in 2022^1^. TB is classically considered “cured” once infection is sterilized. However, recent work has brought attention to the fact that many TB survivors will go on to have permanent sequelae. In fact, up to 50% of the morbidity and mortality associated with TB infection is estimated to accrue after microbiologic cure^2^. Post-TB lung disease (PTLD) is the dominant contributor to this increased morbidity and mortality^3-9^.

While consensus definitions of PTLD are still under active discussion^10^, PTLD has been described to be composed of three contributing pathologies: cavitation, bronchiectasis, and fibrosis^11^. On a histopathologic level, matrix destruction and pathologic remodeling are at the core of PTLD. Suggestive evidence supports the concept that dysregulated matrix destruction underlies the parenchymal cavitation and bronchiectasis seen in PTLD^12-14^; the fibrosis literature points to dysregulated matrix remodeling as underlying fibrogenesis. The specific cells and molecules that contribute to tissue injury and pathologic matrix remodeling in the context of TB infection remain largely unknown.

To date, therapy for pulmonary tuberculosis has been limited to antibiotics aimed at sterilizing infection. Adjunctive host-directed therapy (HDT) has been proposed as a path to either hastening sterilization or improving outcomes after treatment^15^. Adjunctive immunomodulators are in fact routinely used in two types of TB disease: infection of the brain and meninges (tuberculosis meningitis) and infection of the serosal surface of the heart (tuberculosis pericarditis). In those two cases, the addition of immunomodulating steroids to standard antibiotics has been shown to improve functional outcomes^16,17^. In theory, adjunctive use of targeted therapies that reduce pathologic matrix destruction or reduce fibrogenesis together with standard antibiotics could improve functional outcomes after treatment for pulmonary TB. However, our current knowledge of the pathogenesis of tissue damage and remodeling is insufficient to enable the rational selection of candidate adjunctive agents. One significant barrier to a mechanistic understanding of the pathogenesis of distinct components of PTLD is the lack of a small animal model that can be used for hypothesis-generation, hypothesis testing, and pre-clinical studies.

To enable mechanistic and preclinical studies of the fibrotic component of PTLD, we have adapted a standard mouse model of TB infection. Histopathologic patterns of fibrosis identified in tissues from individuals with TB were similar to those seen in Mtb-infected mice. Fibrotic changes were quantifiable and predominantly present in regions of cellular infiltration; fibrosis did not resolve with antibiotic treatment. Inflammation and inflammatory infiltrates resolved slowly, with the dominant inflammatory signatures associated with infection remaining elevated weeks into treatment. A transcriptional signature previously identified as associated with pro-fibrotic macrophages in a range of disorders was upregulated in both carrier- and antibiotic-treated infected lungs.

## Results

### The histopathologic spectrum of fibrotic lesions in human disease

As a starting point for evaluating a potential mouse model, we first sought to characterize the range of fibrotic lesions present in the context of human TB infection. To accomplish this goal, we took advantage of a large human lung tissue repository we had developed in Durban, South Africa comprising surgical resection and post-mortem samples that were collectively representative of the clinical continuum of TB disease (i.e. latent, sub-clinical, active and healed TB), potentially relevant confounders (i.e. anti-TB drug susceptibility, variable HIV co-infection, smoking history) and other important comparators (i.e. normal/healthy lung tissue and known or idiopathic causes of human pulmonary fibrosis). We identified several distinct morphologies of TB-associated fibrosis in human lungs which we then described using a novel, variable combination of useful parameters, including basic lung micro-anatomy (i.e. pleural, interstitial and alveolar fibrosis), TB-induced or TB-related lesional alterations (i.e. fibro-necrotic granulomas and peri-cavitary fibrosis), histopathological severity of disease (i.e. mild, moderate and severe), and chronology of cellular events (i.e. from early fibroblast activity and collagen deposition without architectural distortion to late alveolar replacement-type fibrosis with parenchymal scarring). Fibronecrotic granulomas were identified in the parenchymal and subpleural regions of the lung (**Fig. 1A-C**). In some regions, the necrosis had progressed to cavitation with pericavitary fibrosis (**Fig. 1B**). We identified lesional fibrosis of varying age within the TB granuloma as well as the surrounding parenchyma independent of the necrosis (**Fig. 1D**). Chronic fibro-inflammatory alterations in the lung also showed temporal heterogeneity with early alveolar replacement fibrosis (**Fig. 1E**), early and intermediate interstitial fibrosis (**Fig. 1F-G**), intermediate and late subpleural fibrosis (**Fig. 1F-G**), and late-stage fibrosis, with highly organized, mature collagen and minimal residual cellularity in keeping with scar tissue (**Fig. H**). This micro-anatomical and histomorphological characterization of the spatial distribution and temporal evolution of TB fibrosis in human specimens offered a foundation for the development of a comparative mouse model.

**Figure 1.**
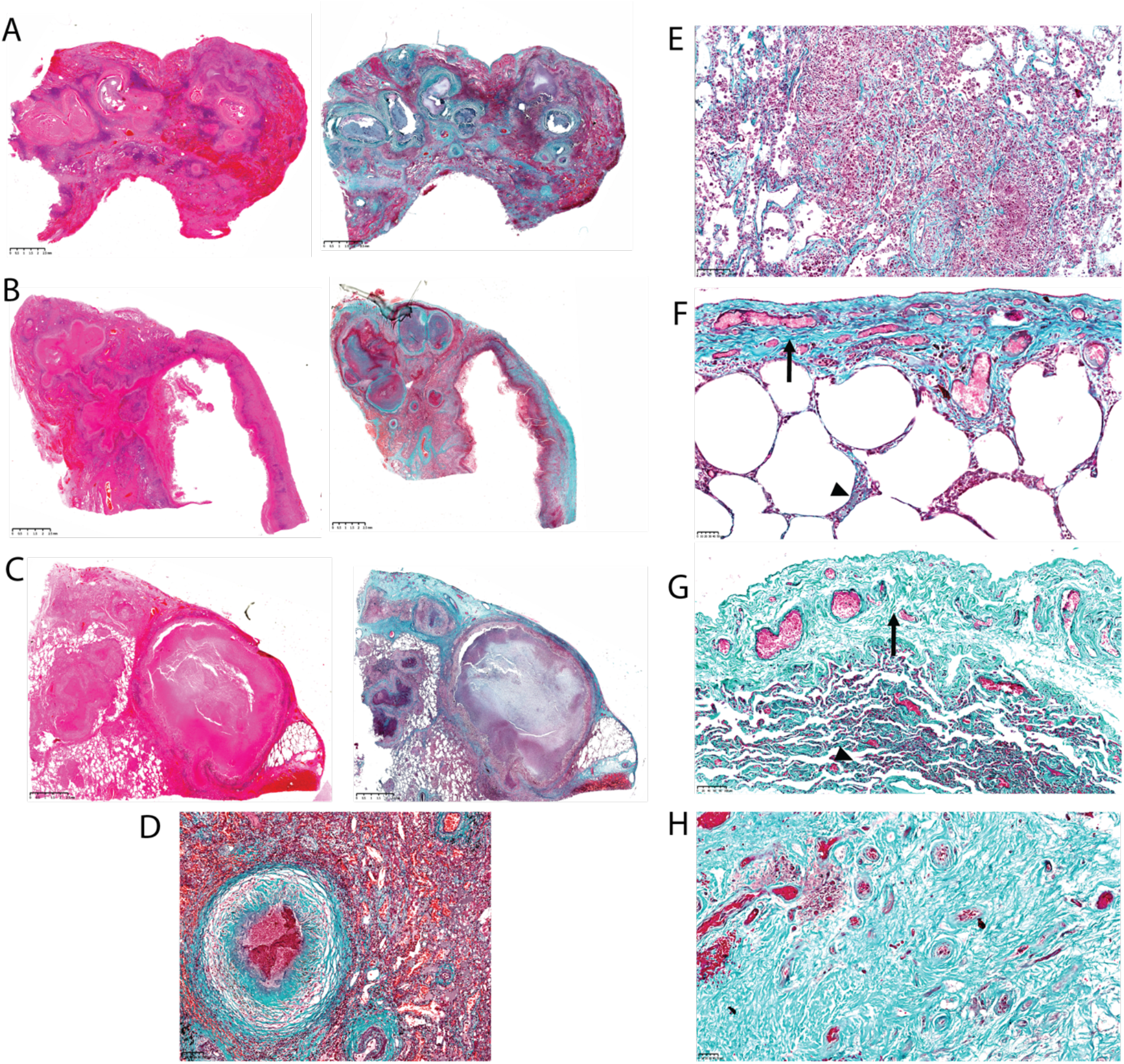
Morphologies of TB-associated fibrosis in human disease. Samples from a human lung tissue repository were stained with Haematoxylin and Eosin (A-C left) or Masson’s Trichrome (A-C right and D-H). (A-C) Multiple necrotizing granulomas in the subpleural and parenchymal regions with variable granuloma-associated fibrosis and surrounding intra-alveolar haemorrhage. (B) includes a conspicuous cavity with adjacent peri-cavitary and pleural fibrosis. (D) Fibrosis of varying age within a necrotizing granuloma as well as the adjacent interstitium and perivascular regions. (E) Fibro-inflammatory alterations, including early alveolar replacement fibrosis with intra-alveolar chronic inflammatory cells and macrophages. (F) Early interstitial fibrosis (arrowhead) and intermediate pleural fibrosis (arrow). (G) intermediate interstitial fibrosis (arrowhead) and late pleural fibrosis (arrow). (H) Late-stage parenchymal scarring/fibrosis effacing normal lung architecture.

### The histopathologic spectrum of fibrotic lesions in a mouse model of TB infection

To develop a quantifiable model for TB-associated lung fibrosis, we used the C3HeB/FeJ model of TB infection^18-20^. Different mouse strains have somewhat distinct courses of disease following infection with Mtb, presenting with a range of bacterial burdens and times to death^21,22^. The C3HeB/FeJ model recapitulates some key aspects of human histopathology, including the development of necrotic lesions with fibrous caps^18-20^. Because of these similarities, this mouse strain has been used extensively for pre-clinical studies. We selected this strain of mouse to investigate the development of pulmonary fibrosis post-treatment.

In accordance with the established model^20^, mice were infected with approximately 100 colony forming units (CFU) of Mtb Erdman via the aerosol route. Bacterial burden plateaued by 3 weeks post-infection, consistent with the timing of the onset of adaptive immunity (**Fig. 2A)**. Notably, cellular infiltrates in the lungs were very modest at the time that bacterial burden plateaued; however, inflammatory infiltrates continued to expand over time (**Fig. 2B-C**). The separation of the plateau in bacterial growth phenotype with progression of pulmonary inflammatory lesions suggested that lung inflammation progresses independent of total bacterial burden and raised the possibility that tissue damage arises from an overly exuberant inflammatory response rather than a direct effect of bacterial effectors^23^.

**Figure 2.**
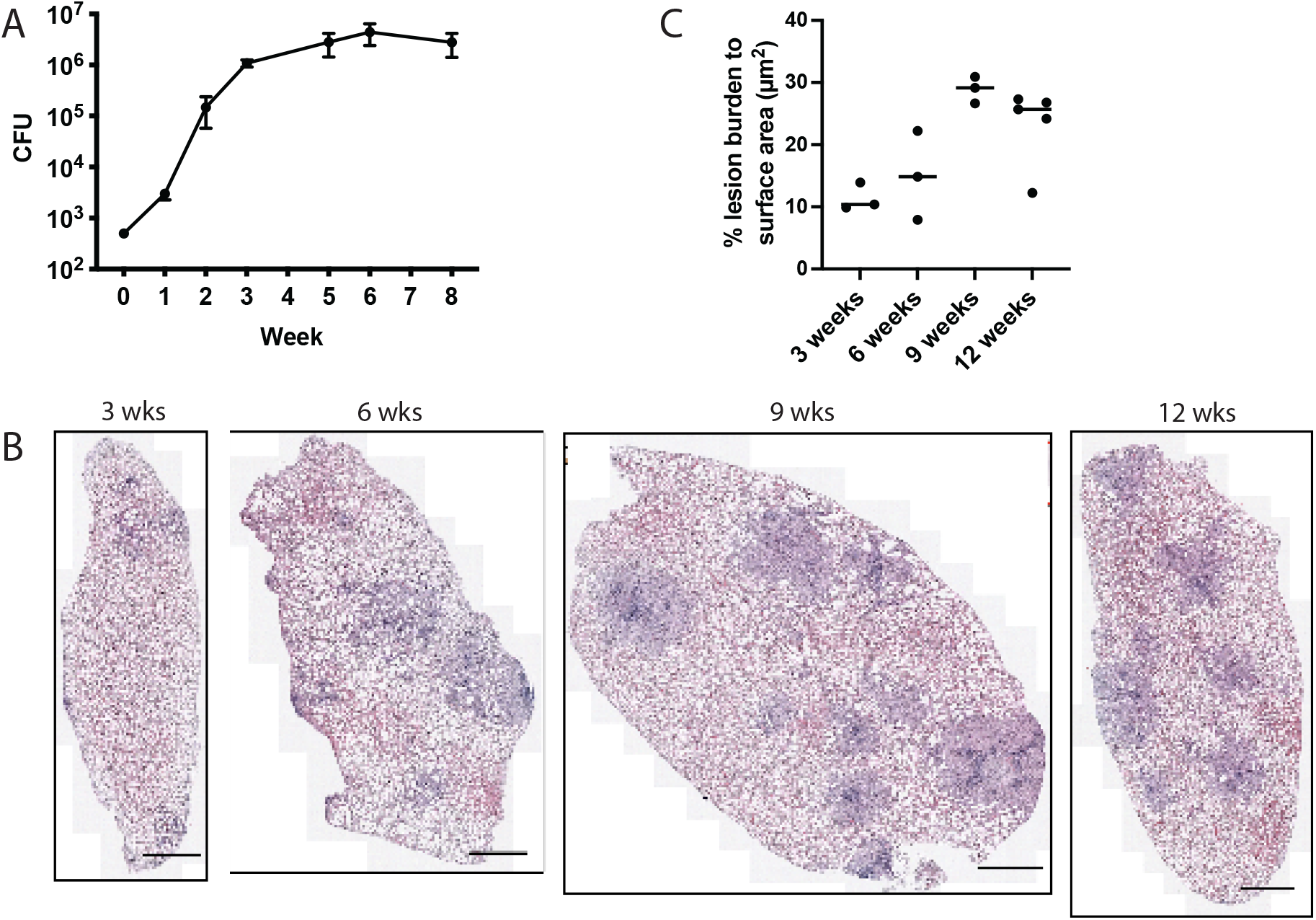
Progression of Mtb infection in the C3HeB/FeJ mouse model. C3HeB/FeJ mice were infected with Mtb Erdman via low dose aerosol. (A) Mice were harvested at the indicated timepoints post-infection and lungs were plated for CFU. (B) Lungs were harvested and formalin-fixed at the indicated timepoints post-infection. Lungs were stained by Hematoxylin and eosin to visualize inflammatory infiltrates. Black bar: 1mm. (C) Quantification of inflammatory infiltrate area based on H&E stained sections.

We next sought to assess whether fibrotic lesions appeared in this mouse model, and if so whether they were similar to lesions in human lung. Using Masson’s Trichrome staining to visualize collagen fibers, we found thickened, linearized collagen in alveolar walls by 9 weeks post-infection, suggestive of early interstitial fibrosis (**Fig. 3A**). Similar to human disease, the histopathology seen in C3HeB/FeJ mice was heterogeneous, with significant mouse-to-mouse and within-mouse variability. By 16 and 20 weeks post-infection, we identified a range of fibrotic histopathologic lesions, including subpleural fibrosis (**Fig. 3B**), early fibronecrotic lesions (**Fig. 3C**), and alveolar replacement fibrosis (**Fig. 3D**). While generally earlier in stage that those observed in human specimens, these lesions suggested that C3HeB/FeJ mice develop histopathologic lesions similar to fibrotic lesions seen in the context of human TB (**Fig. 1**). We thus considered that C3HeB/FeJ mice offer a viable model for studying the pathogenesis of TB-associated fibrosis.

**Figure 3.**
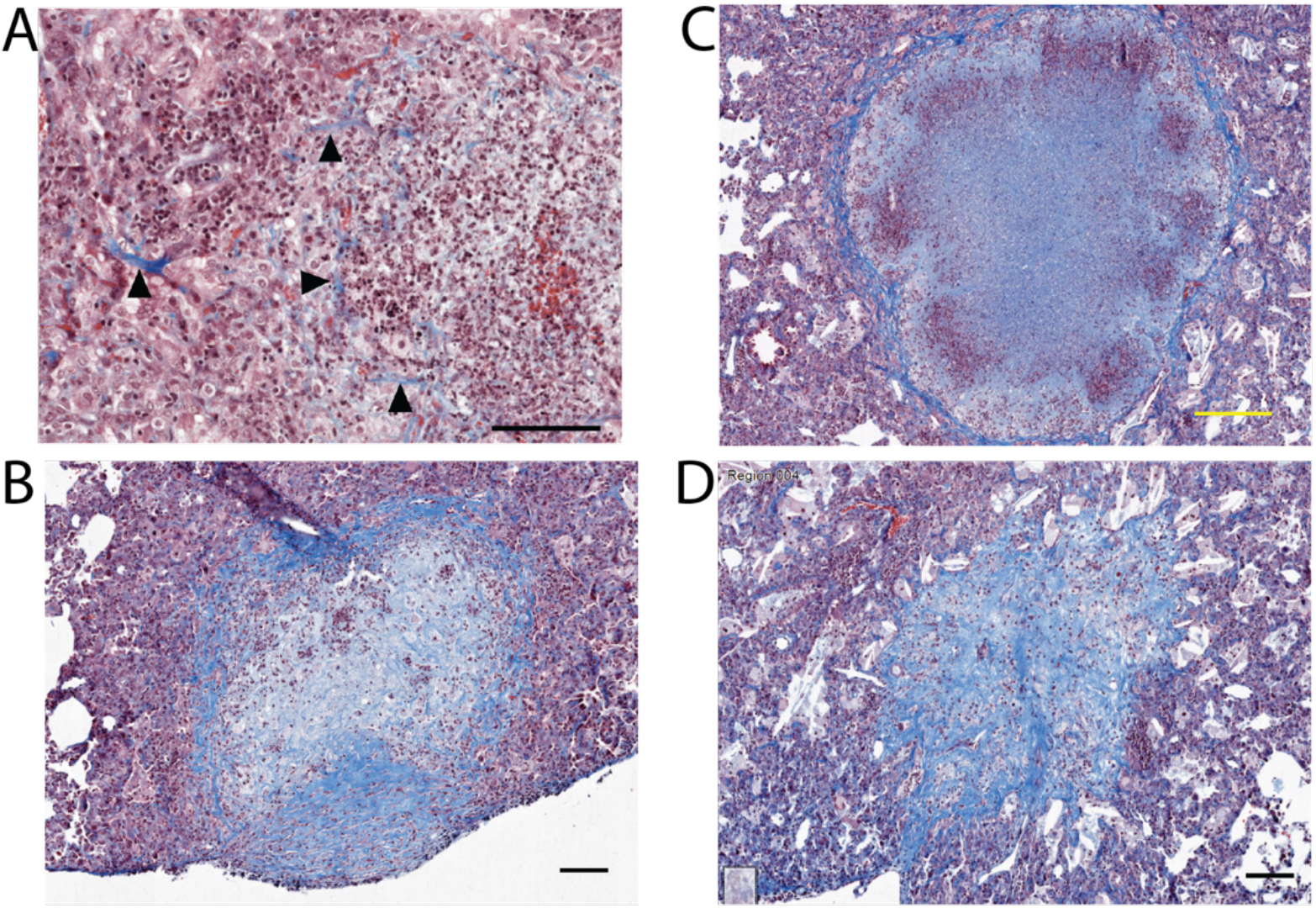
Mtb-infected C3HeB/FeJ mice develop fibrosis in histopathologic patterns similar to human disease. C3HeB/FeJ mice were infected with Mtb Erdman via low dose aerosol. Lungs were harvested at the indicated timepoints post-infection and stained with Masson’s Trichrome to visualize collagen. (A) 9 weeks post-infection, thickened/linearized collagen consistent with early interstitial fibrosis (arrowheads) (B) 12 weeks, subpleural fibronecrotic lesion (C) 14 weeks, necrotizing granuloma with fibrous cap (D) 14 weeks, alveolar replacement fibrosis. Black bars: 100µm. Yellow bar: 200µm.

### Interstitial fibrosis is quantifiable and occurs in regions of cellular infiltration

For mechanistic and pre-clinical studies, we needed a method for quantify fibrosis in our Mtb-infected mouse lungs. To achieve this aim, we deployed the imaging approach of second harmonic generation (SHG) microscopy. SHG microscopy is a label-free nonlinear optical microscopic imaging technique that is highly specific and sensitive to non-centrosymmetric fibrillar collagen and has been used for imaging fibrillar collagen in many tissue types^24-27^. Our Masson’s Trichrome stains suggested that each of the identified types of fibrosis occurred exclusively in regions of inflammatory infiltrates (**Fig. 3**). We thus applied SHG microscopy to specifically quantify fibrillar collagen deposition in regions of inflammatory infiltrates in our Mtb-infected mouse lungs (**Fig. 4A**). By visual inspection, lungs from Mtb-infected mice had significantly more fibrillar collagen deposition than lungs from uninfected mice (**Fig. 4B**). Using pixel density to quantify fibrillar collagen in each image, we found that by 12 weeks post-infection infected mice had significantly more fibrillar collagen than uninfected mice (**Fig. 4C**).

**Figure 4.**
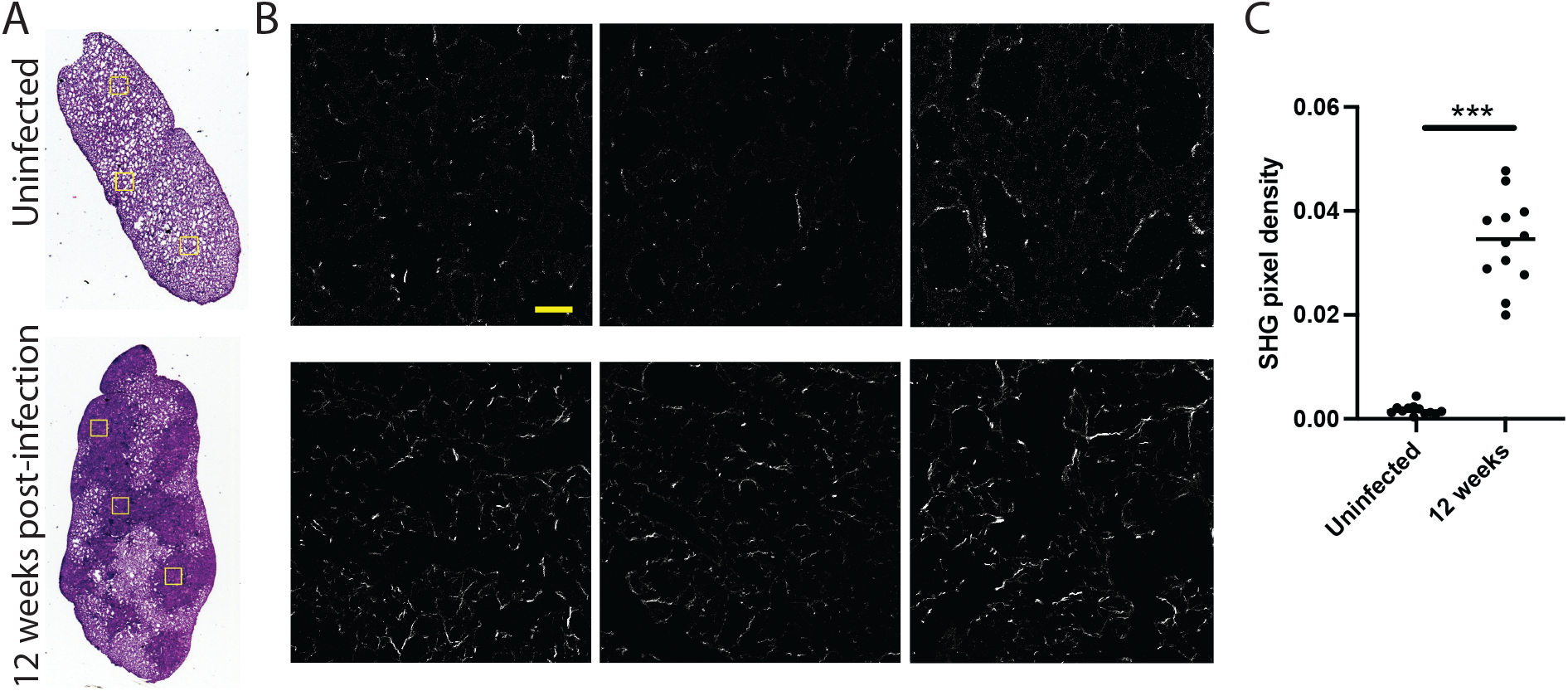
Fibrillar collagen in Mtb-infected mouse lungs is quantifiable by Second Harmonic Generation microscopy. **(A)** Bright-field images for an uninfected lung (top) and a Mtb-infected lung (12-weeks post-infection). (B) SHG images of the regions of interest (ROIs) highlighted in yellow boxes in (A) with the top row showing the uninfected ROIs and the bottom row showing the infected ROIs. (C) Collagen density in 12 ROIs for 4 uninfected samples and 4 infected samples. For visualization purpose, the format of images in (B) was changed from the original 16-bit to 8-bit, and the image contrast in (A) and (B) was adjusted by applying an “Auto” Brightness/Contrast adjustment using Fiji ^46^. *** p-value < 0.0001, Mann-Whitney U test. The ROI size is 410µm by 410 µm. Scale bar equals 50µm.

### Fibrotic changes persist during antibiotic treatment

In human TB, radiographic findings of fibrosis do not typically resolve after treatment. However, in mouse models of some pulmonary fibrotic processes and post-influenza lung fibrosis, fibrosis does improve or resolve entirely^28,29^. To assess whether TB-associated fibrosis persists or resolves in mice after treatment, we next assessed the stability of fibrosis through treatment with standard anti-TB antibiotics. Mice were infected with Mtb Erdman via low-dose aerosol; infection was allowed to progress for 12 weeks. At that time, mice were treated with a standard cocktail of first-line TB drugs, including rifampin (R), isoniazid (I), pyrazinamide (P), and ethambutol (E) via oral gavage (**Fig. 5A**)^30^. Bacterial burden progressively decreased after 2, 4, and 8 weeks of RIPE therapy (**Fig.5B**); the percent of the lung impacted by inflammatory infiltrates decreased with treatment (**Fig. 5C**). Masson’s Trichrome staining demonstrated that although regions of cellularity began to resolve, the histopathologic features of fibrosis, including fibronecrotic granulomas, subpleural fibrosis, and alveolar replacement fibrosis (**Figs. 5D-E**), did not resolve with treatment. Consistent with the persistent fibrosis noted after 2, 4, and 8 weeks of RIPE therapy on Masson’s Trichrome staining, SHG microscopy analysis of fibrosis demonstrated persistent fibrillar collagen in tissue (**Fig. 5F**). These results demonstrate that, consistent with human disease, TB-associated fibrosis in our mouse model does not resolve with antibiotics alone.

**Figure 5.**
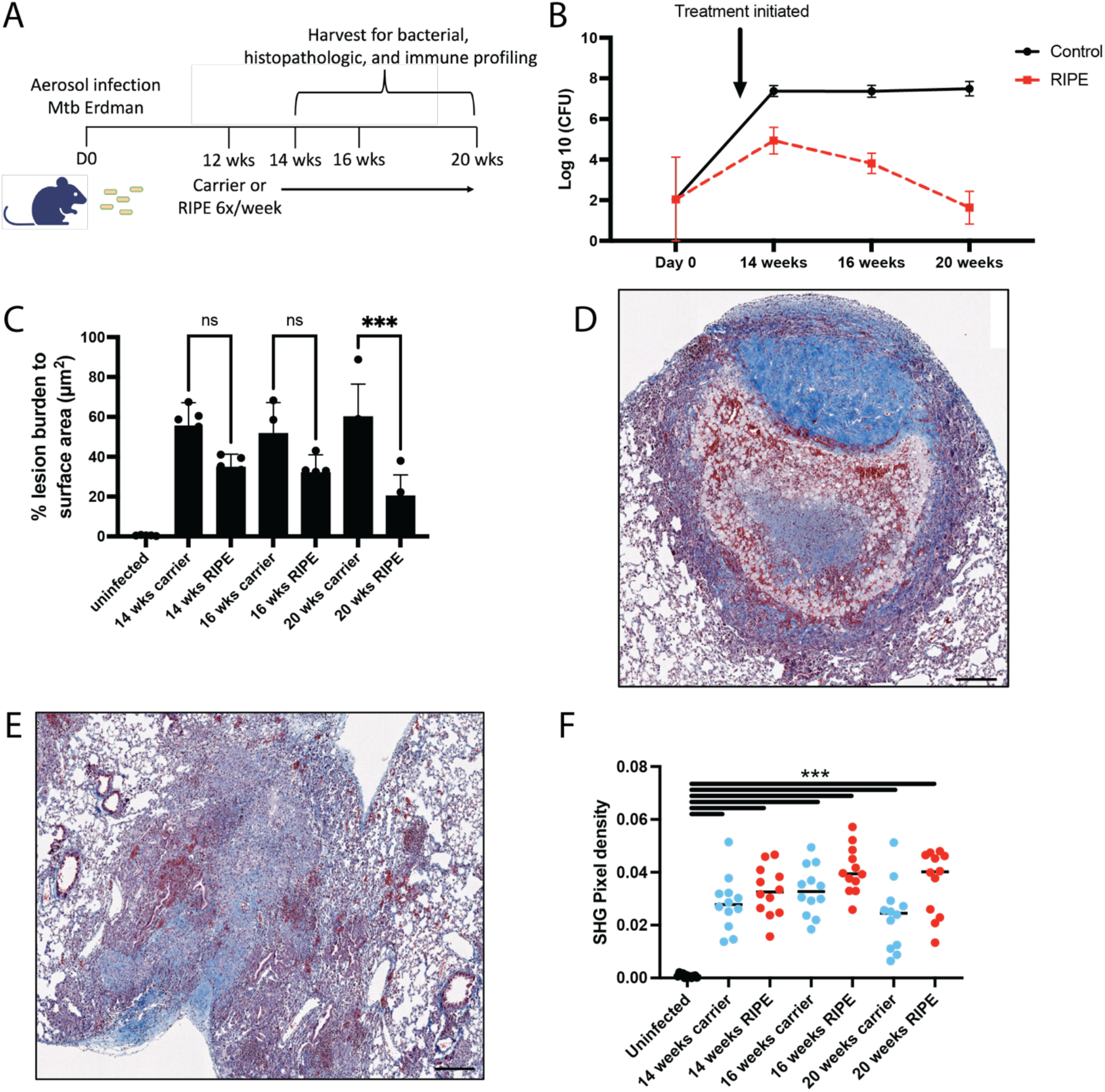
TB-associated fibrosis in mouse lung does not resolve with antibiotic treatment. C3HeB/FeJ mice were infected with Mtb Erdman via low dose aerosol. Infection was allowed to progress for 12 weeks; mice were then treated 6 times per week with RIPE therapy or water carrier control. (A) Infection, treatment, and harvest schema. (B) Lungs were harvested at the indicated timepoints and plated for bacterial burden. (C) Lungs were fixed at the indicated timepoints and stained with H&E to visualize inflammatory infiltrates. QuPath was used to quantify inflammatory infiltrates relative to total lung area. (D-E) Fixed lungs were stained with Masson’s Trichrome to visualize collagen. (D) 16 weeks, subpleural fibronecrotic granuloma (E) 20 weeks, alveolar replacement fibrosis Black bar: 200µm (F) SHG microscopy quantification of fibrillar collagen in carrier- and RIPE-treated mice over time. *** p-value < 0.0001, Mann-Whitney U test.

### Matrix remodeling continues through antibiotic treatment

Longitudinal studies of lung function after completion of treatment for TB suggests that lung function continues to change even after antibiotic treatment is completed^31^. Seeking to assess whether changes occur on a tissue level after treatment initiation, we asked whether matrix remodeling continues after starting anti-TB antibiotics in our mouse model. We used bulk transcriptional profiling on whole mouse lungs to broadly assess the impact of antibiotic treatment on matrix remodeling enzymes (as in **Fig. 5A**). At 14 weeks post-infection, we found that transcript of six matrix metalloproteinases (MMPs) were significantly elevated above baseline expression (**Fig. 6A**). Among the upregulated genes were *Mmp8*, a neutrophil collagenase, *Mmp14*, a collagenase implicated in lung matrix damage early in the course of influenza infection and previously identified in human TB granulomas^13,32^, and *Mmp12*, an elastase noted to be elevated in alveolar macrophages from patients with active TB infection^33^. Strikingly, even after 8 weeks of treatment with antibiotics, expression of most of the six MMPs did not return to baseline levels (**Fig. 6A**). These results suggest that matrix remodeling continues during antibiotic treatment.

**Figure 6.**
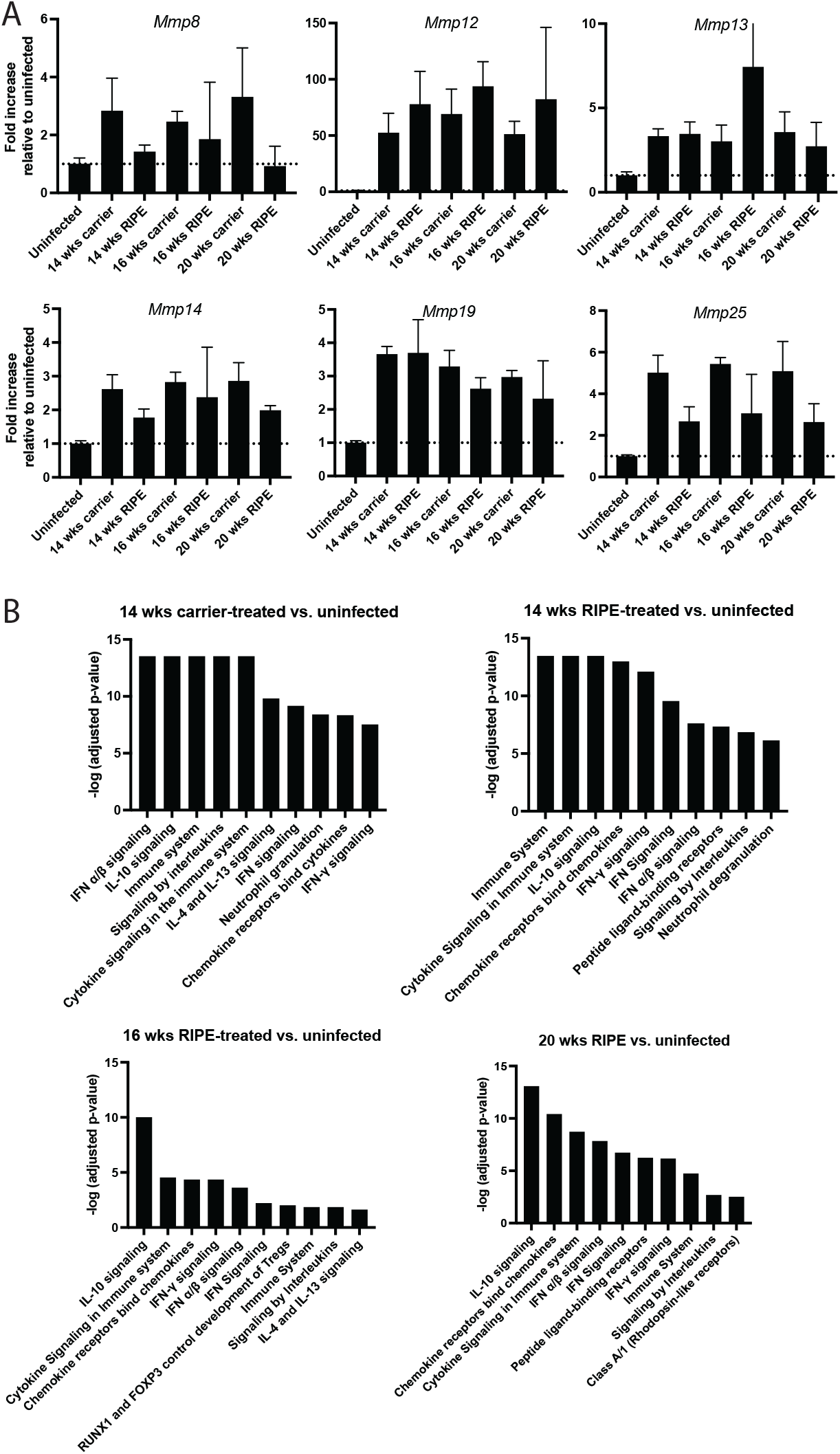
Matrix metalloproteinases and inflammatory gene signatures remain upregulated in lungs during antibiotic treatment for TB. Mice were infected with Mtb Erdman via low dose aerosol and treated as shown in Figure 5A. Lungs were harvested at the indicated timepoints; RNA was isolated from whole lung and comprehensive transcriptomic profiling was performed. (A) Fold increase FPKM of individual MMPs upregulated in the context of TB infection. (B) Differentially expressed genes for each indicated comparison were identified. Reactome Pathway Analysis^36^ was used to analyze genes upregulated more than 2 fold for each comparison shown. The same inflammatory pathways remained significant differentially expressed even after 2, 4, and 8 weeks of RIPE therapy.

### Inflammation resolves slowly during antibiotic treatment

The precise role of the immune response and inflammation in driving TB-associated fibrosis is uncertain. In clinical disease, scarring and fibrosis occurs in regions of prior inflammation. In our mouse model, histopathologically we find that fibrillar collagen deposition occurs in regions of inflammatory infiltrates (**Figs. 4A-B**). If inflammation contributes to acute tissue injury and/or pathologic tissue remodeling in TB-associated fibrosis, ongoing inflammation during and after treatment would be expected to contribute to ongoing remodeling. Evidence from human studies and NHP models in fact suggests that inflammation begins to decrease but does not quickly resolve with antibiotics^34,35^.

We used bulk transcriptional profiling to characterize the impact of antibiotics on the inflammatory state in our mouse model (schematic as in **Fig. 5A**). At 14 weeks post-infection, 1071 genes were upregulated at least 2-fold in infected, carrier-treated mice relative to uninfected mice. As expected, pathway analysis^36^ was consistent with upregulation of innate and adaptive immune responses, including type I and type II IFNs, neutrophil granulation, and IL-10 signaling (**Fig. 6B**). After 2 weeks, 4 weeks, and 8 weeks of RIPE anti-TB therapy, innate and inflammatory gene signatures remained highly upregulated, with 514, 1010, and 431 genes more than 2-fold upregulated in infected, RIPE-treated mice relative to uninfected mice at those timepoints, respectively. Pathway analysis^36^ demonstrated that the same signaling pathways remained significantly upregulated, including IL-10 signaling and type I and type II IFNs (**Fig. 6B**). These results suggest that inflammatory signaling in the lung does not resolve quickly with antibiotics but persists even after weeks of treatment that has largely cleared the burden of live Mtb from the lungs.

### Inflammatory macrophages and macrophage-associated fibrotic transcriptional signatures persist during antibiotic treatment

Macrophages have predominantly been studied as a replicative niche for Mtb in the course of infection. The role that macrophages might play in long-term tissue pathology has not been given substantial consideration to date. However, macrophages are increasingly appreciated to play a role in other forms of pulmonary fibrosis, including the canonical model of pulmonary fibrosis, IPF^37-39^. We considered whether macrophage subpopulations might contribute to fibrogenesis in TB-infected mice. In parallel with transcriptional analysis, we performed immunophenotyping cells in the lungs of RIPE-treated and carrier-treated mice by flow cytometry, with a particular focus on macrophage subpopulations (**Fig. 5A**). At 14 weeks post-infection, infected mice treated with either carrier or RIPE therapy demonstrated a significant reduction in alveolar macrophage populations relative to uninfected controls (**Fig. 7A**). Alveolar macrophages began to repopulate at 16 and 20 weeks post-infection (4 and 8 weeks after initiation of RIPE therapy). Interstitial macrophages and Ly6c^hi^ macrophage populations were increased in infected, carrier-treated mice at 14 weeks post-infection; interstitial macrophages had not returned to baseline levels after 4 and 8 weeks of RIPE therapy (16 and 20 weeks post-infection). Ly6c^hi^ macrophages were elevated at 14 weeks post-infection and had trended down after 4 and 8 weeks of RIPE therapy. These results demonstrate that inflammatory populations of macrophages persist for weeks after initiation of antibiotic treatment in Mtb-infected mouse lungs.

**Figure 7.**
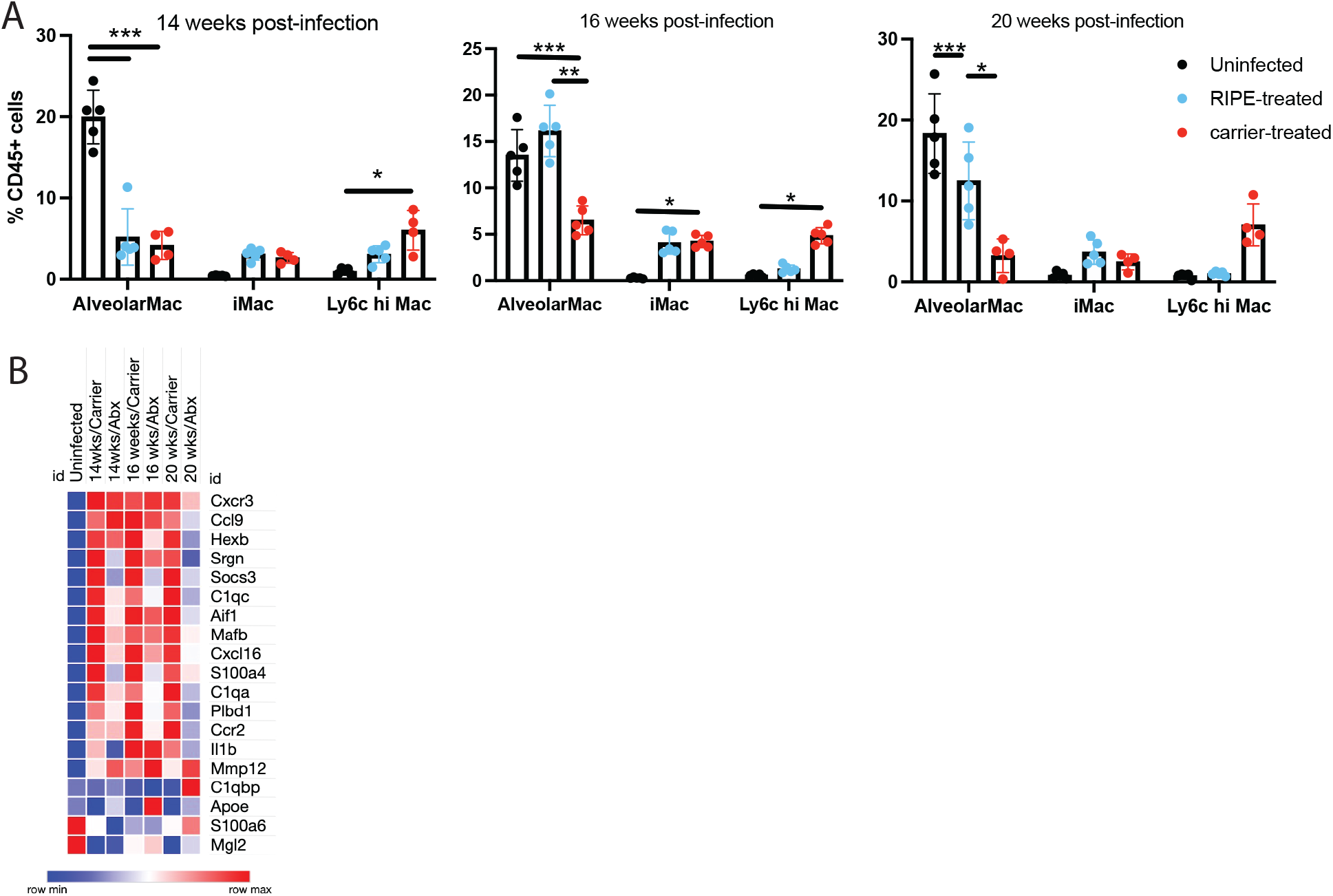
Inflammatory macrophages and pro-fibrotic transcriptional signatures persist through antibiotic treatment. Mice were infected with Mtb Erdman via low dose aerosol and treated as shown in Figure 5A. Lungs were harvested at the indicated timepoints. (A) Single cell suspensions were generated and lungs were antibody stained to identify immune cell subsets^47^. Gating was performed as previously published^45^. (B) RNAseq data from Figure 6 was probed for a gene signature previously identified as associated with transitional macrophages in distinct fibrotic pathologies^39^. Abx indicates RIPE treatment.

The mouse model of IPF has enabled mechanistic investigations of individual cell types that contribute to fibrosis in this process. Recent work coupling single cell-RNAseq with immunofluorescence analysis in this model identified a transitional macrophage associated with fibrosis^39^. Probing our RNAseq dataset for expression of this transcriptional signature, we found that the signature was markedly upregulated at 14 weeks post-infection; it remined upregulated in RIPE-treated mice after 4 weeks of treatment and had not fully resolved to baseline levels after 8 weeks of treatment (**Fig. 7B**). These results suggest that a pro-fibrotic transcriptional signature previously associated with transitional macrophages in distinct fibrotic pathologies is present during TB infection in lung and persists through anti-TB antibiotic treatment.

## Discussion

With millions of individuals treated for TB each year and a significant proportion developing PTLD, TB infection is arguably the single leading cause of pulmonary fibrosis in the world. Adjunctive therapies that could direct healing away from pathologic remodeling toward healthy remodeling could have a substantial impact on global lung health. Lack of a fundamental understanding of fibrogenesis in the context of Mtb infection and the absence of a small animal model have been significant first-order barriers to developing such therapies. Here, we have adapted a mouse model of TB infection for dedicated study of TB-associated lung fibrosis. Benchmarking our results against TB-associated fibrosis from human specimens, we find similar patterns of fibrosis, although the patterns are typically in earlier stages of progression in the mouse model. We anticipate this model and the capacity to quantify fibrosis in the model will form the foundation for future mechanistic and preclinical studies that are strictly necessary for the development future adjunctive therapeutics that can improve outcomes after treatment.

One previous study in mice has raised the question of whether targeting the fibrosis-associated growth factor TGF-β with the IPF drug pirfenidone during treatment for TB infection may actually worsen tissue pathology^40^. While this work demonstrates the importance of caution in approaching potential therapeutic targets, a full mechanistic understanding of the kinetics and molecular and cellular contributors to fibrogenesis in TB infection will identify a broad range of potential therapeutic targets. Ideally, each candidate can then be directly compared for both impact on fibrosis and infection outcomes using small molecule and/or genetic approaches. Some targets are likely to offer more benefit in the context of TB-associated fibrosis than others, and the timing of adding a precision anti-fibrotic agent may be critical for outcome.

One critical question in considering the possibility of adjunctive therapies to improve functional outcomes after TB treatment is whether there is a therapeutic window during which lung health can be preserved or improved. Evidence from the clinical literature in fact suggests that changes in lung function continue through and after antibiotic treatment, providing just that window^31^. Our transcriptional data demonstrates that multiple MMPs remain upregulated in the lungs through weeks of antibiotic treatment, supporting the notion that matrix remodeling is ongoing during this time. Inflammation is likely a major contributor to both ongoing injury and pathologic tissue remodeling in TB infection. Our data similarly support the idea that inflammation continues throughout the course of treatment. Modulating individual matrix remodeling enzymes, the inflammatory milieu during healing, or both holds significant promise for improving functional outcomes and longevity for the millions of individuals treated for TB each year.

## Methods

### Human cohort/ethics statement

The acquisition of ante-mortem and post-mortem human biological samples for scientific research by the Africa Health Research Institute (AHRI) in Durban, South Africa was authorized by the University of KwaZulu-Natal Biomedical Research Ethics Committee (class approval study number: BCA535/16). The surgical resection lung tissue samples in the cohort were predominantly from patients undergoing pneumonectomies or lobectomies for TB at King Dinizulu Hospital Complex, a tertiary center for TB patients in Durban (study identifier: BE019/13). Written informed consent was obtained from every study participant. These patients included HIV-positive and HIV-negative individuals with varying smoking histories and co-morbidities. The major indication/s for surgery were advanced fibrocavitary disease, bronchiectasis, recurrent pulmonary infections, haemoptysis and/or failed pharmacological therapy for drug-sensitive and drug-resistant strains of *Mtb*. Post-mortem TB lung samples in the repository were mainly from decedents who were reportedly asymptomatic and demised from causes other than TB (i.e. latent and/or subclinical TB). Notably, the autopsy samples also included healthy/normal and neoplastic lung tissue for comparison as well as other known or idiopathic causes of pulmonary fibrosis. The overall sample cohort for our histopathological appraisal of TB-associated lung fibrosis is therefore collectively representative of the complex clinical continuum of TB disease (i.e. latent, sub-clinical, active and healed TB), with/out HIV infection and varying anti-TB drug-susceptibility profiles (i.e drug-sensitive, multi-drug resistant and extreme drug-resistant *Mtb* stains).

### Human tissue histopathology

For haematoxylin & eosin (H&E) staining, samples of lung were aseptically removed and fixed in 10% neutral buffered formalin. Samples were processed in a vacuum filtration tissue processor using a xylene-free method with isopropanol as the main substitute fixative. Tissue sections were then embedded in paraffin wax. Sections were cut at 4 µm, baked at 60°C for 15 min, dewaxed through two changes of xylene and rehydrated through descending grades of alcohol to water. Routine H&E staining was performed whereby slides were placed in haematoxylin for 5 minutes, washed in tap water for 2 minutes, blued in lithium carbonate for 1 minute, rinsed in tap water for 2 minutes, and counter stained with eosin for 5 minutes before a final rinse in tap water for 2 minutes. Slides were dehydrated in ascending grades of alcohol, cleared in xylene, and mounted in Distyrene, Plasticizer, and Xylene (DPX).

For Masson’s Trichrome (MT) staining, samples of lung tissue were cut into 3 µm sections, mounted on charged slides, and heated at 56°C for 15 minutes. Mounted sections were dewaxed in xylene followed by rinsing in 100% ethanol and one change of 95% ethanol. Slides were then washed under running water for 2 minutes. Next, slides were treated with Bouin’s fixative for 30 minutes in a 60°C oven. After fixation, slides were allowed to cool for 10 minutes and then washed in running tap water until sections appeared clear. Fixed slides were then stained with celestine blue for 5 minutes, followed by rinsing in running tap water. Slides were stained in Mayer’s hematoxylin for 5 minutes and rinsed again in running tap water. To blue the tissue sections, slides were immersed in lithium carbonate for 1 minute, then washed once more in running tap water. Slides were then stained with biebrich scarlet for 3 minutes, rinsed in running tap water before being treated with phosphotungstic acid for 15 minutes or until the biebrich scarlet no longer diffused. Finally, slides were washed in running tap water, then counterstained with light green for 1 minute. Sections were blot dried. The counterstained step was repeated to achieve the desired result. Slides were then dehydrated in ascending grades of alcohol, cleared in xylene, and mounted in Distyrene, Plasticizer, and DPX.

### Bacterial strains and mouse infections

Mouse infections were carried out as previously published^41^. In brief, the Mtb Erdman was grown to mid-log phase in 7H9 media supplemented with 0.05% tween-20, 0.2% glycerol, and 10% Middlebrook OADC. Bacteria were washed twice in PBS, then used in aerosol infections using an AeroMP device (Biaera) with a goal of 100 CFU implanted per mouse. All mouse work was performed according to protocols approved by the Massachusetts General Hospital Institutional Animal Care and Use Committee.

### Antibiotic treatments

Mice were treated in accordance with published protocols for standard anti-TB treatment^30^. In brief, mice were treated via oral gavage with a suspension of rifampin (10mg/kg), isoniazid (10mg/kg), pyrazinamide (150mg/kg) and Ethambutol (100mg/kg) six times pe week. Control mice were gavaged with an equal volume of water six times per week.

### Mouse Lung Histopathology

Mouse lungs were fixed in 10% formalin. Paraffin embedding, H&E staining, and MT staining of lung sections were performed by the Massachusetts General Hospital Rodent Histopathology Core.

### Second Harmonic Generation Microscopy

A custom-built SHG microscope was used to detect the collagen of the lung tissue samples. The principle of SHG and an early version of this SHG system have been described in detail in previous work^24^. Briefly, in this study, the SHG system was built on an upright microscope Olympus BX61 (Olympus, Tokyo, Japan). A Chameleon Ultra Ti:Sapphire laser (Coherent Inc, Santa Clara, CA) was used to deliver light with a wavelength of 895nm to the sample using a 20X/0.75NA air immersion objective (Olympus, Tokyo, Japan). SHG signal from the collagen was filtered with a bandpass filter 445/20nm (Semrock, Rochester, NY) in the forward channel 1. The autofluorescence signal from the sample was not filtered in the forward channel 2. The signals from both channels were detected using two identical photon counting GaAsP photomultiplier tubes H7421-40 (Hamamatsu, Hamamatsu, Japan). Circular polarization was implemented such that collagen fibers in all orientations were excited equally. Three representative regions of infiltrates were imaged from each sample. All images were collected at 512 pixels by 512 pixels resolution (1 pixel = 0.8015µm) with consistent acquisition settings using in-house developed open-source laser-scanning acquisition software OpenScan (https://github.com/openscan-lsm/OpenScan). The images were then stitched together for each region of interest using the Fiji Grid/Collection Stitching plugin ^42^. An established collagen open-source quantification tool “CurveAlign” (v5.0, https://github.com/uw-loci/curvelets/releases/tag/5.0)^43^ was used to calculate the collagen density in the infiltration regions. Based on the pair of autofluorescence image and brightfield image, three representative regions with the size of 512 pixels x 512 pixels (or 410 um x 410 um) were annotated by avoiding undesirable SHG signals from e.g. blood vessels in each selected infiltration regions in the autofluorescence image. The annotations were then applied to the corresponding co-registered SHG image. The collagen density was calculated as the number of collagen pixels (defined as the pixel whose SHG intensity is larger than 5) divided by the total number of pixels in the region of interest.

### Quantification of granuloma area

Digital images of the hematoxylin and eosin-stained slides were obtained with the TissueFAXS SQL Confocal Slide Scanner. Quantitative analysis was performed blinded to treatment allocation and slides pertaining to the mice, as previously published^44^. In brief, for each treatment group, a total of 3-4 scanned slides were reviewed. The lung lesion burden to whole surface area was quantified using the open-source software QuPath (https://qupath.github.io/), as described [CT]. The lung inflammatory lesions to whole surface area were quantified and multiplied by 100 to obtain the percentage.

### RNAseq analysis

Mouse lungs were soaked in RNAlater and frozen at −80 until the completion of the experiment. RNA was then isolated using Trizol (Invitrogen) according to the manufacturer’s protocols. RNAseq libraries were generated and sequenced by the Dana Farber Molecular Biology Core Facility. RNAseq data was then aligned using STAR method. Fold change in expression with respect to the uninfected mice was evaluated by DESeq2. The samples were then clustering by applying Likelihood-ratio test (LRT) with a threshold of log2Foldchange ≥ 1.5 and p_adj_< 0.01 in R. Heatmap for the gene signatures associated with the fibrotic lung was created using MORPHEUS algorithm (https://github.com/cmap/morpheus.R) in R.

### Immunophenotyping by flow cytometry

Immunophenotyping of murine lungs was performed as previously described^45^. Briefly, freshly harvested murine lungs were dissociated in a GentleMACS Dissociator (Miltenyi Biotec) in digestion buffer (RPMI with 10mM HEPES, DNAse I 50ug/ml, Liberase TM 100ug/ml, and 2% FBS) using the m_lung_01 program first followed by a 30-minute incubation at 37C and then the m_lung_02 program. After filtration with a 70-uM filter, samples were treated with RBC Lysis Buffer (Sigma-Aldrich) for 5min. The lysis step was stopped by addition of FACS buffer (PBS with 2% FBS and 2 mM EDTA) and the samples were washed once before staining. Cells were first incubated with fixable viability dye eFluor 455UV (Invitrogen), followed by Fc receptors block (TruStain FcX, clone 93, BioLegend) and then stained with a panel of immunophenotyping antibodies (clone, dilution, manufacturer) at room temperature for 30min: CD45 BUV395 (30-F11, 1:400, BD Biosciences), CD24 BV510 (M1/69, 1:500, BioLegend), I-A/I-E Pacific Blue (M5/114.15.2, 1:1200, BioLegend), CD64 Pe/Cyanine7 (X54-5/7.1, 1:50, BioLegend), CD11c PerCP (N418, 1:200, BioLegend), CD11b (M1/70, 1:1500, BioLegend), Ly-6G BV605 (1A8, 1:1500, BioLegend), Ly-6C AF700 (HK1.4, 1:300, BioLegend) and SiglecF PE-CF594 (E50-2440, 1:1000, BD Biosciences). After the incubation, the samples were washed in PBS, then fixed with 4% paraformaldehyde (Santa Cruz Biotechnology), and filtered through a 70µm filter (BD biosciences) before acquisition. Data was acquired on a BD Symphony flow cytometer (BD Biosciences) using BD FACSDiva software (BD Biosciences) and analyzed using FlowJo software (v10.7.1, BD). Macrophage subpopulations are expressed as percentage out of live CD45+ cells.

## Acknowledgements

The authors gratefully acknowledge funding from NIH grants R01CA238191 (KWE), P41GM135019 (KWE and BAC), R21AI146813 (AKB), R33AI138280 (AJCS), and the Wellcome Leap Delta Tissue Program (AJCS). Work performed in the Ragon Institute BSL3 Core and the Dana Farber Molecular Biology Core Facility was supported in part by the Harvard University Center for AIDS Research (P30 AI060354). The authors would like to thank Drs. Roi Avraham, Inna Solomonov, Irit Sagi, Sara Auld, Greg Bisson, Rachel Knipe, Benjamin Medoff, and Ms. Cathleen Krabak for helpful discussions.

